# Sustained anti-obesity effects of life-style change and anti-inflammatory interventions after conditional inactivation of the activin receptor ALK7

**DOI:** 10.1101/2020.07.28.224881

**Authors:** Raj Kamal Srivastava, Ee-Soo Lee, Eunice Sim, New Chih Sheng, Carlos F. Ibáñez

**Affiliations:** Department of Physiology, National University of Singapore, Singapore 117597, Singapore; Life Sciences Institute, National University of Singapore, Singapore 117456, Singapore; Department of Neuroscience, Karolinska Institute, Stockholm 17177, Sweden

**Author notes:** Present address: Indira Gandhi National Tribal University, Amarkantak, MP 484887, India.

**Keywords:** activin, adipose tissue, adrenergic signaling, C/EBPα, diet-induced obesity, Na-salicylate

## Abstract

Life-style change and anti-inflammatory interventions have only transient effects in obesity. It is not clear how benefits obtained by these treatments can be maintained longer term, specially during sustained high caloric intake. Constitutive ablation of the activin receptor ALK7 in adipose tissue enhances catecholamine signaling and lipolysis in adipocytes, and protects mice from diet-induced obesity. Here, we investigated the consequences of conditional ALK7 ablation in adipocytes of adult mice with pre-existing obesity. Although ALK7 deletion had little effect on its own, it synergized strongly with a transient switch to low-fat diet (life-style change) or anti-inflammatory treatment (Na-salicylate), resulting in enhanced lipolysis, increased energy expenditure, and reduced adipose tissue mass and body weight gain, even under sustained high caloric intake. By themselves, diet-switch and salicylate had only a temporary effect on weight gain. Mechanistically, combination of ALK7 ablation with either treatment strongly enhanced the levels of β3-AR, the main adrenergic receptor for catecholamine stimulation of lipolysis, and C/EBPα, an upstream regulator of β3-AR expression. These results suggest that inhibition of ALK7 can be combined with simple interventions to produce longer-lasting benefits in obesity.

## Introduction

Life-style change is arguably the safest and most prevalent intervention used to counteract obesity. However, as difficult as it is to adhere to a severe diet and exercise regime, even more difficult it is to maintain the lost weight over longer periods of time, particularly after the end of a weight-lowering regime (van Baak and Mariman, 2019). A study of participants undergoing an intensive diet and exercise intervention as part of “The Biggest Loser” televised weight loss competition found that on average 70% of the lost weight was regained 6 years after the competition (Fothergill et al., 2016). Weight loss results in a slowing of resting metabolic rate, a metabolic adaptation that acts to counter weight loss, and a likely contributing factor to weight regain (Müller and Bosy-Westphal, 2013). Interventions that increase energy expenditure may reduce the risk of weight regain after a life-style change, and help to stabilize the lower weight in the post-regime period. In addition, obesity has been associated with a state of inflammation in adipose tissue that contributes to development of insulin resistance and diabetes (Hotamisligil, 2006). Simple anti-inflammatory interventions, such as salicylates and its derivatives, have shown efficacy at reducing adipose tissue inflammation and improving insulin sensitivity and glucose tolerance in both rodents and humans (Fleischman et al., 2008; Yuan et al., 2001). Despite these effects, salicylate treatment does not significantly affect the BMI (body mass index) of obese individuals or body weight in mouse models of obesity (Adapala et al., 2012; Anderson et al., 2014). Anti-inflammatory treatments may however result in reduced fat mass and weight loss when combined with interventions that increase energy expenditure, as suggested by the dual effects of physical exercise on body weight and adipose tissue inflammation (Krüger et al., 2016).

ALK7 is a receptor with intrinsic serine-threonine kinase activity that mediates the effects of a subset of ligands from the Transforming Growth Factor-Beta (TGF-β) superfamily, including the adipokines activin B and growth and differentiation factor 3 (GDF-3) (Andersson et al., 2008; Bertolino et al., 2008; Rydén et al., 1996). In complex with the type II TGF-β superfamily receptor ActRIIB, ligand binding activates ALK7 to promote phosphorylation of Smad2 and Smad3 (Jörnvall et al., 2001; Rydén et al., 1996), key mediators of canonical nuclear signaling by members of the TGF-β superfamily. ALK7 is highly expressed in differentiated adipocytes from both rodent and human adipose tissues (Andersson et al., 2008; Carlsson et al., 2009; Kang and Reddi, 1996; Murakami et al., 2012). Our previous studies have shown that ALK7 is dispensable for mouse development (Jörnvall et al., 2004), but mice that lack ALK7 in adipose tissue have enhanced catecholamine signaling in adipocytes, increased lipolysis and energy expenditure, and, accordingly, show resistance to diet-induced obesity (Guo et al., 2014). Thus, ALK7 normal function in adipocytes is to facilitate fat accumulation by suppressing the expression and signaling of adrenergic receptors, thereby contributing to catecholamine resistance in obesity (Guo et al., 2014). In addition, experiments using a chemical-genetic approach showed that acute inhibition of ALK7 kinase activity in adult mice concomitant with a switch to high-fat diet protected the mice from diet-induced obesity (Guo et al., 2014), indicating that ALK7 can be inhibited acutely and does not need to be eliminated from birth to affect obesity. Recent studies have identified polymorphic variants in the human *Acvr1c* gene (encoding ALK7) which affect body fat distribution and protect from type II diabetes (Emdin et al., 2019; Justice et al., 2019). These findings indicate that ALK7 has similar functions in humans as in rodents, and suggest that strategies to suppress ALK7 signaling may be beneficial to combat human obesity. However, studies in animal models that deleted or inactivated ALK7 prior to diet-induced obesity left open the question of whether ablation of ALK7 may have comparable effects in mice with pre-existing obesity, which is the situation normally encountered clinically.

In this study we used a genetic approach to conditionally inactivate ALK7 in adult obese mice following 12 weeks on high fat diet. We combined conditional ALK7 ablation with either life-style change (i.e. switch form high-fat to low-fat diet) or anti-inflammatory treatment (Na-salicylate), and assessed effects on weight gain, fat deposition and different biochemical and physiological parameters in several adipose tissue depots.

## Results

### Acute ablation of ALK7 in adipose tissue of obese mice enhances the beneficial effects of life-style change

In order to conditionally inactivate ALK7 in adipose tissue of adult mice we bred *Alk7*^flox/flox^ mice (Guo et al., 2014) to *AdipoQ*^CreERT2^ mice expressing tamoxifen-inducible Cre recombinase under the control of the mouse *Adipoq* locus, encoding adiponectin, an adipokine specifically expressed in fully differentiated adipocytes (Sassmann et al., 2010). In the studies described below, we have compared *Alk7*^flox/flox^ mice (herein referred to as control mice) with *Alk7*^flox/flox^::*AdipoQ*^CreERT2^ mice (herein called mutant mice) after 3 daily consecutive administrations of tamoxifen as indicated in the Methods section. Control and mutant mice subjected to 12 week high fat diet (HFD) were treated with tamoxifen and sacrificed 2 weeks later to assess *Alk7* mRNA expression in adipocytes and stromal vascular fraction (SVF), containing adipocyte precursors as well as immune and vascular cells, from epididymal and subcutaneous (inguinal) white adipose tissue depots (EWAT and SWAT, respectively). In both adipose depots, expression of *Alk7* mRNA was reduced by more than 95% to almost background levels in mature adipocytes, indicating a highly efficient deletion (Supplementary Figure S1A). As expected, no *Alk7* mRNA expression was detected in SVF of either control or mutant mice (Supplementary Figure S1B), as this fraction does not contain mature adipocytes. Expression of the gene encoding ALK4, a receptor related to ALK7, was unaffected in the mutant mice (Supplementary Figure S1C, D).

Loss of ALK7 resulted in a small decrease (≈ 2g) in body weight in mutant mice kept on HFD compared to controls, starting 2 weeks after tamoxifen treatment and maintained for up to 5 weeks (Figure 1A-C, light blue and orange curves). We note that tamoxifen administration to obese mice by itself induced a 3 to 4g loss in body weight in all groups, an effect that has been reported previously (Ye et al., 2015). To assess a possible interaction between inactivation of ALK7 and life-style change interventions, a parallel cohort of HFD-fed control and mutant mice was switched to standard Chow diet (SD) two days after the last tamoxifen administration; all groups were kept for an additional 5 weeks (Figure 1A). Both control and mutant mice showed a steady drop in body weight after the diet switch, but weight loss occurred more rapidly and to a greater extent in the mutant mice, which differed by >5g from controls by one week after diet switch (Figure 1B, C, dark blue and red curves). Food intake was not different between control and mutant mice (Supplementary Figure S2A, B). Compared to control mice, weights of EWAT and SWAT depots in mutant mice were reduced by about 20 to 25% on HFD, but by 45 to 50% after the diet switch (Figure 1D, E). In line with reduced adipose tissue mass, the levels of circulating leptin were greatly reduced in the mutants after diet switch, but were not significantly different between genotypes in the mice kept on HFD (Figure 1F). As expected, plasma insulin levels were reduced after switch to SD, but did not differ between genotypes in either regime (Figure 1G). Glucose tolerance was improved in the mutants under both diet regimes (Figure 1H), while a trend (which did not reach statistical significance) was also observed for insulin sensitivity (Figure 1I). Indirect calorimetry showed elevated O_2_ consumption and CO_2_ production in the mutants under both diet regimes, but significantly more so in the diet switch group (Figure 1J, K). As a result, energy expenditure was also increased in the mutants during both light and dark periods, showing a significant synergy between ablation of ALK7 expression and diet switch (Figure 1L). Respiratory exchange ratio (RER) was lower under constant HFD, reflecting a more prominent use of fat as energy source, but it was not different between genotypes (Figure 1M). Despite elevated energy expenditure in the mutants, locomotor activity was not different between genotypes (Supplementary Figure S3A, B), suggesting increased metabolic rate after ablation of ALK7. Adipocyte size in EWAT and SWAT was significantly reduced in the mutants compared to controls after diet switch, but remained essentially unchanged between genotypes in the cohorts that stayed on HFD (Figure 1N-P). Interestingly, adipose tissue apoptosis, assessed by TUNEL staining, was significantly reduced in the mutants after diet switch compared to all other groups (Figure 1Q-S), indicating that ALK7 ablation and life-style change synergize to reduce adipocyte cell death produced by prolonged high caloric intake.

**Figure 1.**
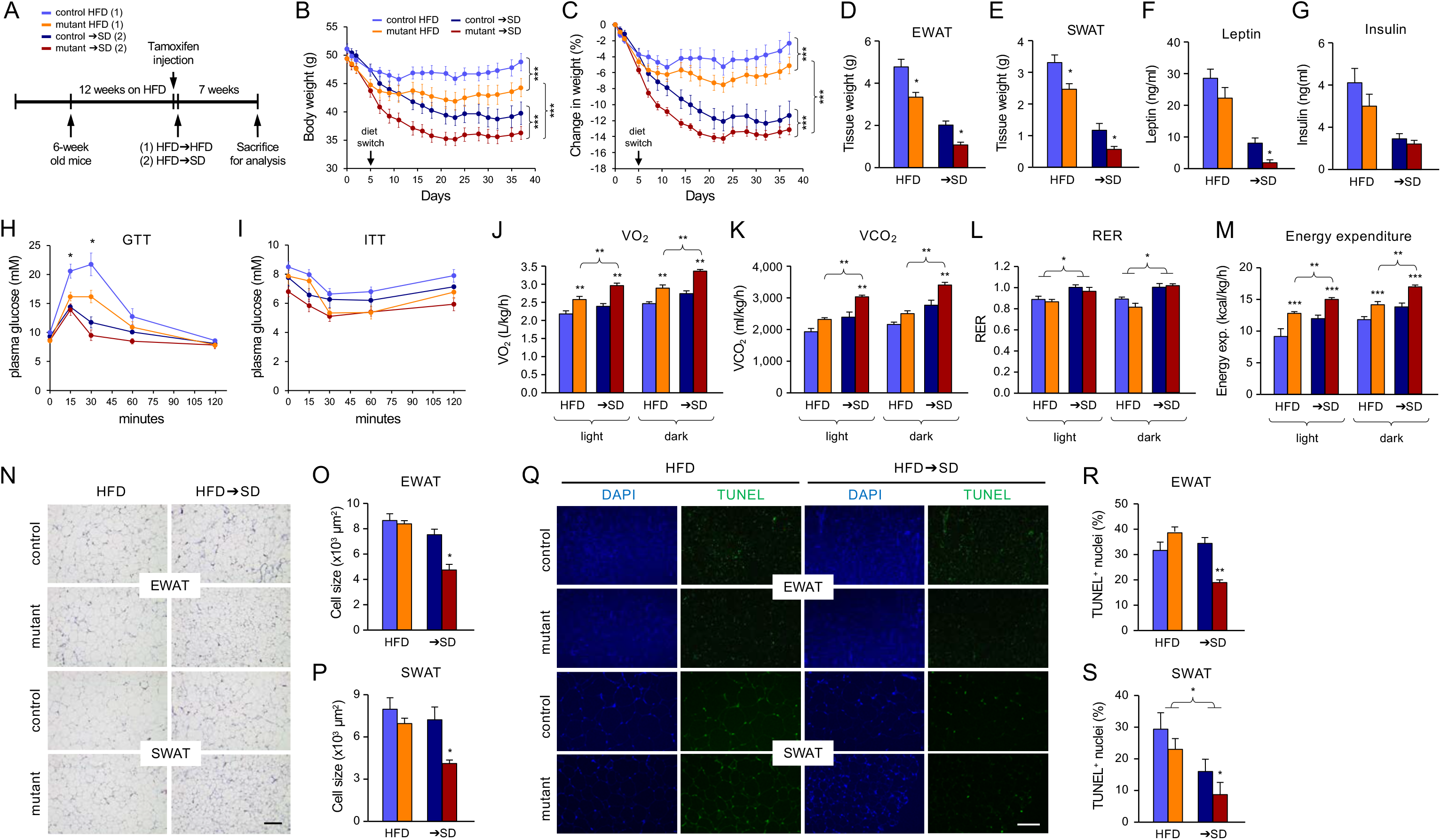
Acute ablation of ALK7 in adipose tissue of obese mice enhances weight loss after life-style change. (A) Schematic of the diet switch protocol. After 12 weeks on HFD, mice were injected with tamoxifen, and one cohort of control and mutant mice (1) was kept in HFD, while the second cohort (2) was switched to SD. Both cohorts were then kept for additional 5 weeks. The color scheme used in all panels of this figure is shown above. (B, C) Body weights of control and mutant mice in constant high-fat diet (HFD) or after a switch to standard diet (→SD) as indicated. Values are expressed as weight in grams (B) or percent change (C) ± SEM. N=6 mice per group. ***, P<0.001; 2-way ANOVA. (D, E) Weights of epididymal (EWAT) and subcutaneous (SWAT) white adipose tissues in control and mutant mice after constant HFD or diet switch (→SD). Shown are mean values ± SEM. N=6 mice per group. *, P≤0.05 vs. control →SD (2-way ANOVA). (F, G) Plasma levels of leptin (EF) and insulin (G) measured by ELISA in control and mutant mice after constant HFD or diet switch (→SD). Shown are mean values ± SEM. N=6 mice per group. *, P<0.05 vs. control →SD (2-way ANOVA). (H, I) Glucose tolerance test (GTT) (H) and Insulin tolerance test (ITT) (I) performed following acute glucose (H) or insulin (I) injection in control and mutant mice after constant HFD or diet switch (→SD). Shown are mean values ± SEM. N=6 mice per group. *, P<0.05 vs. all other groups (2-way ANOVA). (J-L) Oxygen consumption (VO_2)_ (J), CO_2_ production (VCO_2)_ (K), energy expenditure (L) and respiratory exchange ratio (RER) (M) measured by indirect calorimetry at room temperature in control and mutant mice after constant HFD or diet switch (→SD). RER values close to unity indicate carbohydrate usage, while values around 0.7 indicate fat consumption. Shown are mean values ± SEM. N=6 mice per group. *, P<0.05; **, P<0.01; ***, P<0.001 vs. control (within same diet), or vs. HFD (within same genotype), or between diets, as indicated (2-way ANOVA). (N-P) Micrographs of hematoxylin/eosin staining of adipose tissue sections (N) in control and mutant mice after constant HFD or diet switch (→SD). Quantifications of cell size in EWAT (O) and SWAT (P) are expressed as mean ± SEM. N=6 mice per group. *, P<0.05 vs. control (2-way ANOVA). (Q-S) Micrographs of TUNEL staining of adipose tissue sections (Q) in control and mutant mice after constant HFD or diet switch (→SD). Quantifications of % TUNEL positive nuclei in EWAT (R) and SWAT (S) are expressed as mean ± SEM. N=6 mice per group. *, P<0.05; **, P<0.01 vs. control mice or between diets, as indicated (2-way ANOVA).

Next, we assessed the level of expression of a battery of markers of adipose tissue differentiation, function and inflammation in EWAT and SWAT depots of mutants and controls after constant HFD or diet switch. As expected, both depots showed decreased levels of *Leptin* mRNA after switch to SD; interestingly, this reduction was significantly more prominent in the mutants (Figure 2A). One of the more salient differences displayed by the mutants compared to controls in both fat depots and after switch to SD, was the strong induction of mRNAs encoding adrenergic receptors *Adrb1* and *Adrb3* (Figures 2B, C). These changes suggest that ALK7 inactivation and life-style change synergize to restore catecholamine sensitivity in adipose tissue after nutrient overload. In agreement with this, levels of phosphorylated hormone sensitive lipase (HSL), reflecting the activation of a key enzyme for lipolysis downstream of adrenergic signaling, was also significantly elevated in adipose tissue of mutant mice after diet switch (Figures 2D, E). In keeping with elevated adrenergic signaling and P-HSL levels in adipocytes, lipolysis was also elevated in mutants switched to standard diet, both under basal conditions as well as induced by the adrenergic agonist CL316243 (Figures 2F, G). Mechanistically, we found that the level of C/EBPα, a key inducer of adipogenesis and transcriptional regulator of *Adrb3* (Dixon et al., 2001), was differentially increased in the mutants compared to controls (Figures 2H-J). At the protein level, not only did diet switch increased C/EBPα levels, but also synergized very strongly with ALK7 inactivation (Figures 2I, J). Expression of C/EBPα in adipose tissue is actively suppressed by Smad3 (Choy and Derynck, 2003), one of the key mediators of intracellular signaling by TGF-β superfamily receptors, including ALK7. Increased C/EBPα expression in EWAT and SWAT lacking ALK7 is in agreement with this, and parallels the strong induction of *Adrb3* mRNA observed in these tissues. Examination of mRNA expression for a battery of inflammatory markers, including the inflammatory cytokine Interleukin-6 (IL-6), Monocyte chemoattractant protein-1 (Mcp1), which regulates infiltration of macrophages in adipose tissue (Deshmane et al., 2009), and the macrophage markers F4/80, Mrc1, Itgax and Clec10a, revealed a marked reduction of inflammation in adipose tissues after diet switch (Figures 2K-P). For *Mrc1* and *Clec10a* mRNAs, their reduction was most prominent in EWAT than SWAT (Figures 2N, O). In the case of *Mcp1* mRNA, we observed synergistic effects of diet switch and ALK7 ablation (Figure 2L). Together, our analysis of the effects of acute ALK7 deletion in obese mice suggests a strong synergy with life-style change in maintenance of lost body weight, reduced adipose tissue mass, reduced apoptosis, enhanced adrenergic signaling and lipolysis, and reduced adipose tissue inflammation.

**Figure 2.**
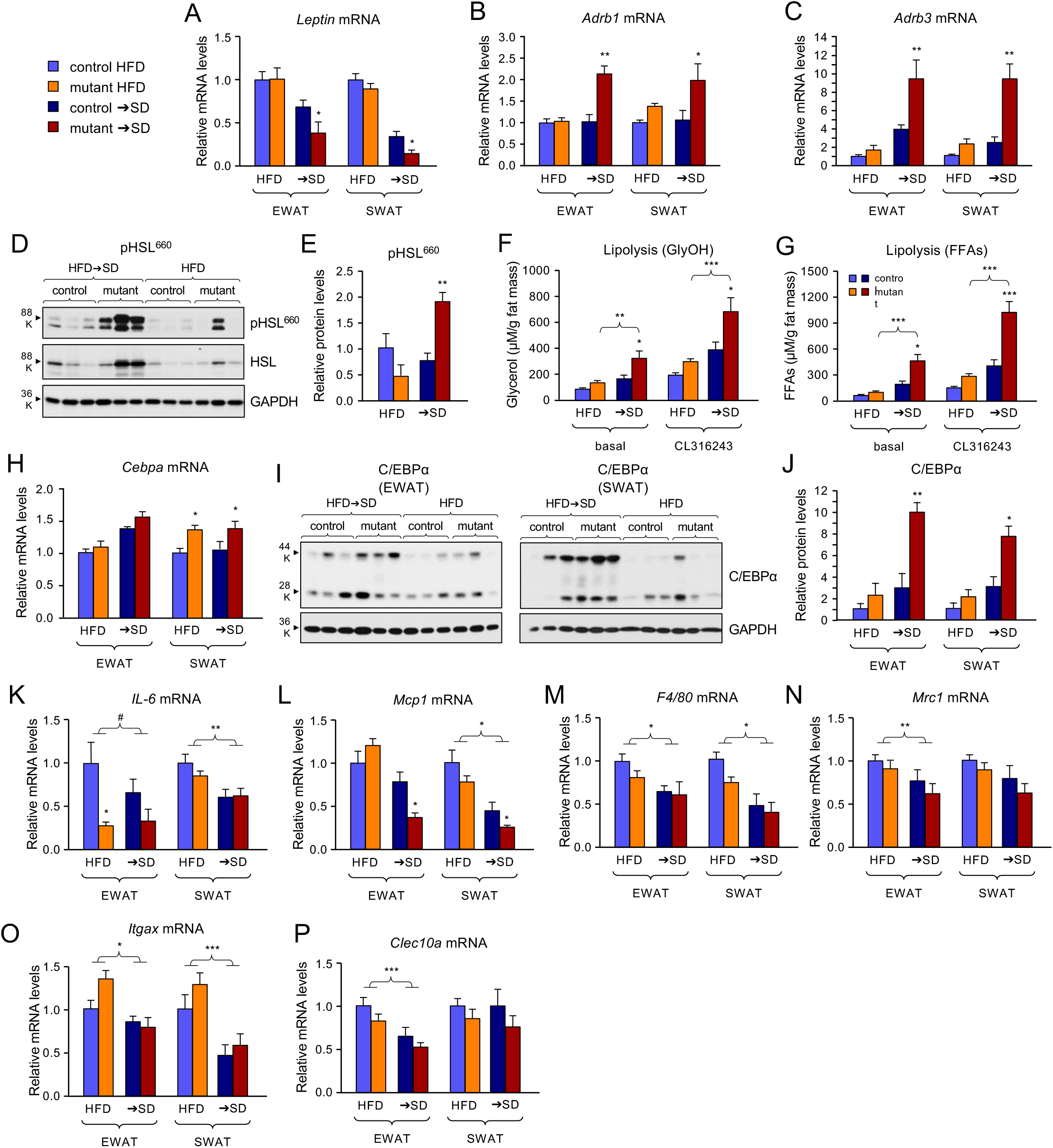
Increased adrenergic signaling and reduced inflammation in adipose tissues of obese mutant mice after switch to standard diet. (A-C) Q-PCR determination of *Leptin* (A), *Adrb1* (B) and *Adrb3* (C) mRNA expression in EWAT and SWAT of control and mutant mice after constant HFD or diet switch (→SD). The values were normalized to mRNA levels in HFD control mice, and are presented as mean ± SEM. N=6 mice per group. *, P<0.05; **, P<0.01 vs. control (within same diet) (2-way ANOVA). The color scheme used in all panels of this figure is shown on the left. (D, E) Western blot analysis of hormone sensitive lipase (HSL) phosphorylated at Ser^660^ (pHSL^660^) in EWAT and SWAT of control and mutant mice after constant HFD or diet switch (→SD). Representative blot shows results of 3 animals per group. Histogram (E) shows quantification of pHSL^660^ levels normalized to total HSL and expressed as mean ± SEM. N=6 mice per group. **, P<0.01 vs. control (within same diet) (2-way ANOVA). (F, G) Assessment of lipolysis as plasma glycerol (F) or free-fatty acids (FFAs) (G) concentration (μM) normalized to fat mass in control and mutant mice after constant HFD or diet switch (→SD), under either basal conditions or following induction by injection of the beta-adrenergic agonist CL316243. Shown are mean values ± SEM. N=6 mice per group. *, P<0.05; **, P<0.01; ***, P<0.001 vs. control (within same diet), or vs. HFD (within same genotype), as indicated (2-way ANOVA). (H) Q-PCR determination of *Cebpa* mRNA expression in EWAT and SWAT of control and mutant mice after constant HFD or diet switch (→SD). The values were normalized to mRNA levels in HFD control mice, and are presented as mean ± SEM. N=6 mice per group. *, P<0.05 vs. control (within same diet) (2-way ANOVA). (I, J) Western blot analysis of C/EBPα protein levels in EWAT and SWAT of control and mutant mice after constant HFD or diet switch (→SD). Representative blots show results of 3 animals per group. Histogram (J) shows quantification of C/EBPα levels normalized to GAPDH and expressed as mean ± SEM. N=6 mice per group. *, P<0.05; **, P<0.01 vs. control (within same diet) (2-way ANOVA). (J-P) Q-PCR determination of *Cebpa* (J), *IL-6* (K), *Mcp1* (L), *F4/80* (M), *Mrc1* (N), *Itgax* (O) and *Clec10a* (P) mRNA expression in EWAT and SWAT of control and mutant mice after constant HFD or diet switch (→SD). The values were normalized to mRNA levels in HFD control mice, and are presented as mean ± SEM. N=6 mice per group. ^#^, P=0.065; *, P<0.05; **, P<0.01; ***, P<0.001 vs. control (within same diet), or between diets, as indicated (2-way ANOVA).

### ALK7 ablation in adipose tissue of obese mice sustains the anti-obesity effects of life-style change after return to high caloric diet

In order to assess whether the beneficial effects of combined ALK7 ablation and life-style change had a longer lasting effect on metabolism, we subjected a separate cohort of mice to a double-switch paradigm, in which obese mice that had been switched to SD went back to HFD for additional 5 weeks (→SD→HFD), and compared them with a group that remained on HFD all the time (Figure 3A). In agreement with the single diet switch study, the effects of 7 weeks switch to SD were more pronounced after ALK7 deletion than in control mice (Figure 3B, C, dark blue and red curves). Interestingly, after these mice were switched back to HFD, control mice gained significantly more weight than mutant mice, even surpassing the body weight of mice that had been kept on HFD all the time (Figure 3B, C). On the other hand, mutant mice regained weight at a much slower pace, and were able to maintain a significant difference compared to the groups that remained under constant HFD (Figure 3B, C). Food intake was not different between control and mutant mice after either diet switch (Supplementary Figure S2C, D). Mutant mice, but not control mice, maintained reduced EWAT and SWAT depots and lower plasma leptin levels after the switch back to HFD (Figure 3D-F), while plasma insulin levels were not changed among groups at the end of the study (Figure 3G). In contrast, double-switched control mice showed higher EWAT deposition and leptin levels (Figure 3D,F), as well as reduced glucose tolerance and insulin sensitivity (Figure 3H, I), than their counterparts kept on constant HFD. Adipose tissue in double-switched mutants showed elevated levels of mRNAs encoding adrenergic receptors (Figure 3J, K) and C/EBPα (Figure 3L). In contrast, the levels of these markers in double switched control mice were not different from those in mice kept in constant HFD (Figure 3J-L). These results suggest that ALK7 ablation helped to sustain the anti-obesity effects of life-style change, even after return to a high caloric diet regime.

**Figure 3.**
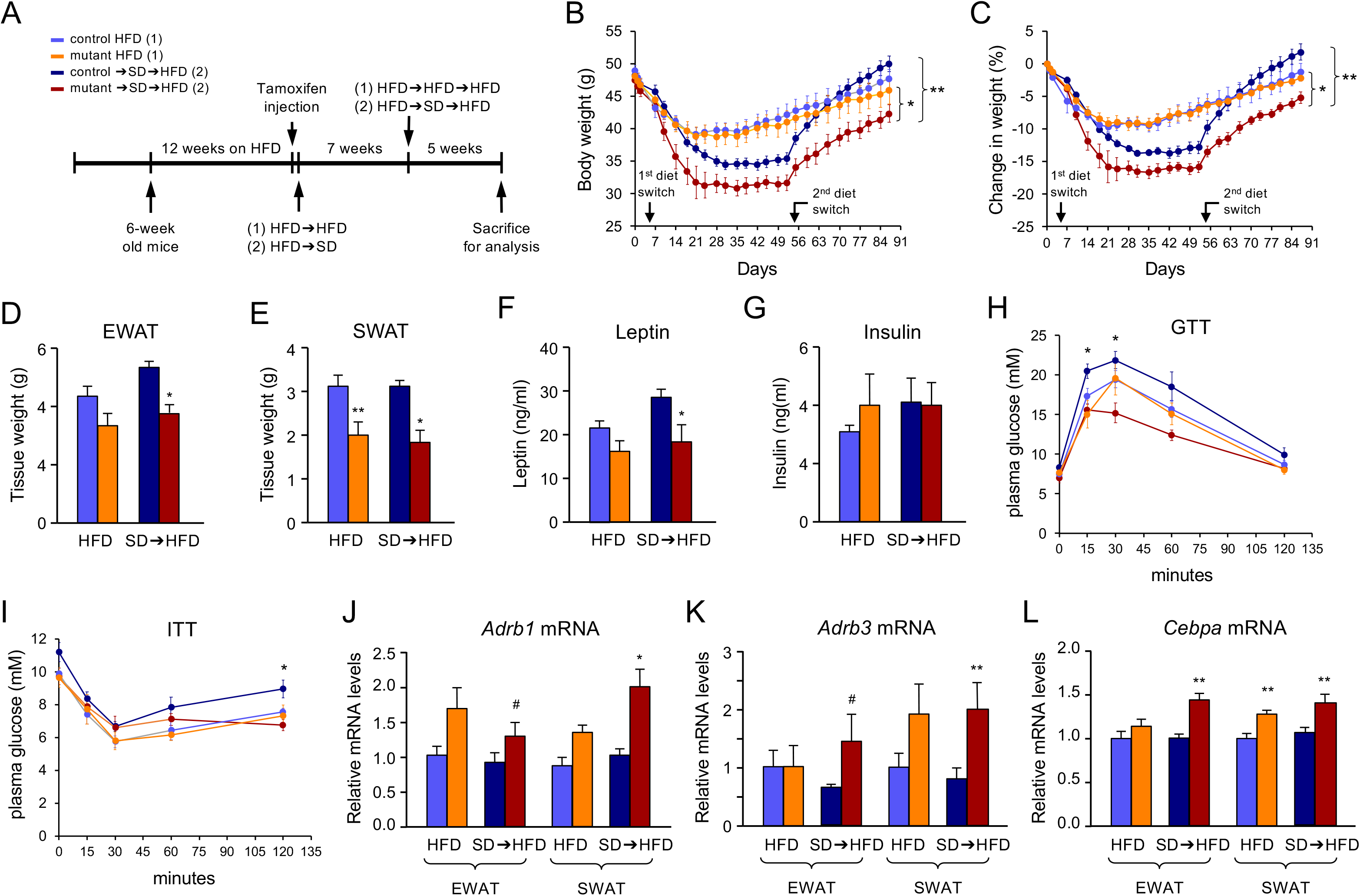
ALK7 ablation in adipose tissue of obese mice sustains the anti-obesity effects of life-style change after return to high caloric diet. (A) Schematic of the double-switch protocol. One cohort of control and mutant mice (1) was maintained in constant HFD. The second cohort (2) was switched to SD after 12 weeks on HFD, then switched back to HFD after 7 weeks, and kept on HFD for additional 5 weeks. The color scheme used in all panels of this figure is shown above. (B, C) Body weights of control and mutant mice in constant high-fat diet (HFD) or after double-switch to standard diet and back to HFD (→SD→HFD) as indicated. Tamoxifen injections started at day 0. Values are expressed as weight in grams (A) or percent change (B) ± SEM. N=6 mice per group. *, P<0.05: **, P<0.01; 2-way ANOVA. (D, E) Weights of epididymal (EWAT) and subcutaneous (SWAT) white adipose tissues in control and mutant mice after constant HFD or double-switch (SD→HFD). Shown are mean values ± SEM. N=6 mice per group. *, P≤0.05 vs. control (within same diet) (2-way ANOVA). (F, G) Plasma levels of leptin (F) and insulin (G) measured by ELISA in control and mutant mice after constant HFD or double-switch (SD→HFD). Shown are mean values ± SEM. N=6 mice per group. *, P<0.05 between control and mutant SD→HFD (2-way ANOVA). (H, I) Glucose tolerance test (GTT) (H) and Insulin tolerance test (ITT) (I) performed following acute glucose (H) or insulin (I) injection in control and mutant mice after constant HFD or double-switch (SD→HFD). Shown are mean values ± SEM. N=6 mice per group. *, P<0.05 vs. All other groups (2-way ANOVA). (J-L) Q-PCR determination of *Adrb1* (J), *Adrb3* (K) and *Cebpa* (L) mRNA expression in EWAT and SWAT of control and mutant mice after constant HFD or double-switch (SD→HFD). Values were normalized to mRNA levels in HFD control mice, and are presented as mean ± SEM. N=6 mice per group. #, P<0.1; *, P<0.05; **, P<0.01 vs. control (within same diet) (2-way ANOVA).

### Acute ablation of ALK7 in adipose tissue synergizes with salicylate to attenuate diet-induced obesity

The synergistic effects of ALK7 deletion and a life-style change intervention underlined the fact that multiple pathways contribute to the profound changes that take place in adipose tissue during obesity. In addition to catecholamine resistance, a hallmark of adipose tissue in obesity is a state of pronounced inflammation, with infiltration of large numbers of immune cells and activation of macrophages which, through secretion of inflammatory cytokines and other messengers, impact on metabolic functions both locally and in distant tissues, such as muscle and liver (Chawla et al., 2011; Reilly and Saltiel, 2017). We reasoned that perhaps combining ALK7 deletion with a simple anti-inflammatory intervention may yield better results than either intervention on its own. We induced ALK7 deletion by tamoxifen injection in mutant and control mice that had been 12 weeks on high-fat diet, starting at the same time treatment with salicylate on one half of mutants and controls. All mice were subsequently kept on high-fat diet for additional 9 weeks (Figure 4A). Food intake was not different between control and mutant mice (Supplementary Figure S2E). Initially, both groups on salicylate showed a 5-6 gram weight loss, but this was transient in control mice, which subsequently gained weight at a higher rate, catching up with the vehicle groups by the 5th week (Figure 4B, C). In contrast, mutant mice on salicylate could maintain reduced weight for several weeks, increasing slowly only after the 5th week. Interestingly, however, they were the only group that ended up with the same weight after 9 weeks on high fat diet; all other mice had gained 5 to 10 grams by the end of the study (Figure 4C). In agreement with this, mutants on salicylate was the only group that showed significantly reduced fat depot mass (Figure 4D, E). In line with previous observations (Fleischman et al., 2008; Yuan et al., 2001), mice on salicylate, both mutants and controls, showed improved glucose tolerance and higher insulin sensitivity than mice on vehicle (Figure 4F, G). However, only mutants showed significant reductions in adipocyte cell size (Figure 4H-J) and apoptosis (Figure 4K-M), particularly when treated with salicylate.

**Figure 4.**
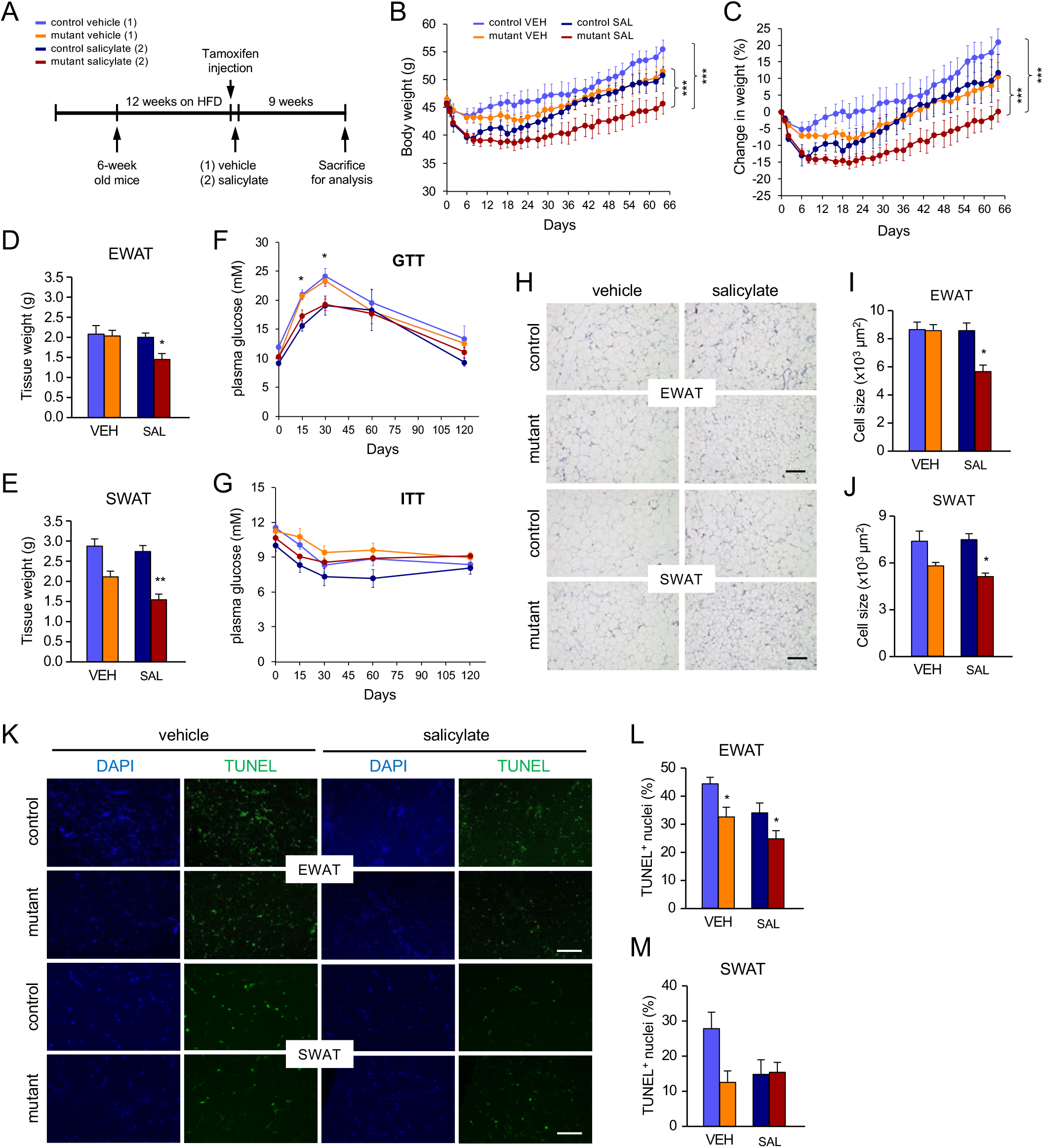
Acute ablation of ALK7 in adipose tissue of obese mice synergizes with salicylate to attenuate diet-induced obesity. (A) Schematic of the salicylate procedure. After 12 weeks on HFD, mice were injected with tamoxifen, and one cohort of control and mutant mice (1) remained on normal drinking water, while the second cohort (2) received drinking water containing salicylate. Both cohorts were then kept for additional 9 weeks on HFD. The color scheme used in all panels of this figure is shown above. (B, C) Body weights of obese control and mutant mice treated with salicylate (SAL) or vehicle (VEH). Tamoxifen injections started at day 0. Values are expressed as weight in grams (A) or percent change (B) ± SEM. N=6 mice per group. ***, P<0.001; 2-way ANOVA. (D, E) Weights of epididymal (EWAT) and subcutaneous (SWAT) white adipose tissues in obese control and mutant mice treated with salicylate (SAL) or vehicle (VEH). Shown are mean values ± SEM. N=6 mice per group. *, P≤0.05; **, P<0.01 vs. control SAL (2-way ANOVA). (F, G) Glucose tolerance test (GTT) (F) and Insulin tolerance test (ITT) (G) performed following acute glucose (G) or insulin (G) injection in control and mutant mice treated with salicylate (SAL) or vehicle (VEH). Shown are mean values ± SEM. N=6 mice per group. *, P<0.05 (between VEH and SAL groups) (2-way ANOVA). (H) Micrographs of hematoxylin/eosin staining of adipose tissue sections in control and mutant mice treated with salicylate (SAL) or vehicle (VEH). (I, J) Quantifications of cell size in EWAT (I) and SWAT (J) are expressed as mean ± SEM. N=6 mice per group. *, P<0.05 vs. control SAL (2-way ANOVA). (K) Micrographs of TUNEL staining of adipose tissue sections in control and mutant mice treated with salicylate (SAL) or vehicle (VEH). (L, M) Quantifications of % TUNEL positive nuclei in EWAT (L) and SWAT (M) are expressed as mean ± SEM. N=6 mice per group. *, P<0.01 (between genotypes) (2-way ANOVA).

Analysis of markers of adrenergic signaling and inflammation further highlighted the basis of the synergy between ALK7 ablation and anti-inflammatory treatment. Adipose tissues of the mutants, under either salicylate or vehicle, showed elevated levels of adrenergic signaling markers, including *Adrb1* and *Adrb3* mRNAs, as well as C/EBPα; intriguingly, these changes were often more pronounced in the mutants treated with salicylate (Figure 5A-D). In agreement with this, lipolysis was also significantly elevated in mutant mice that were treated with salicylate (Figure 5E, F). As expected, several markers of inflammation were reduced in mice treated with salicylate, both mutants and controls, including mRNAs encoding IL-6, TNFa (encoding tumor necrosis factor alpha, the apical regulator of the pro-inflammatory cascade), Mcp1, and macrophage markers F4/80, Mrc1 and Clec10a (Figure 5G-L). Mutants treated with salicylate often showed stronger reduction of inflammatory markers than the other groups. Together, our studies combining salicylate treatment and acute ALK7 deletion in adipose tissue indicate a strong synergy between the two interventions, and support the notion that blunting ALK7 signaling in adipose tissue can be most effective as anti-obesity strategy when inflammation is concomitantly reduced.

**Figure 5.**
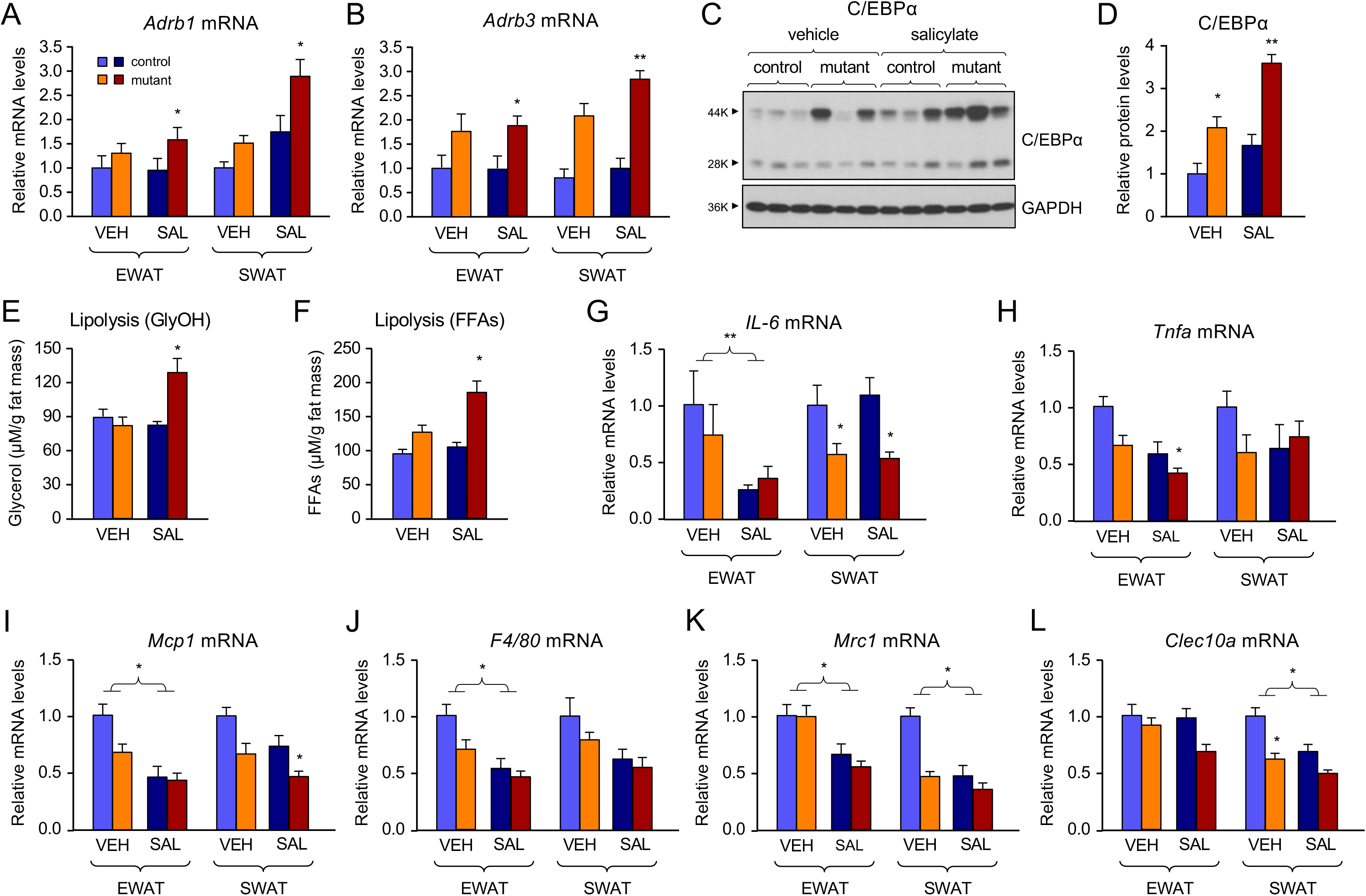
Acute ablation of ALK7 in adipose tissue of obese mice synergizes with salicylate to increase adrenergic signaling and reduce inflammation. (A, B) Q-PCR determination of *Adrb1* (A) and *Adrb3* (B) mRNA expression in EWAT and SWAT of control and mutant mice treated with salicylate (SAL) or vehicle (VEH). The values were normalized to mRNA levels in VEH control mice, and are presented as mean ± SEM. N=6 mice per group. *, P<0.05; **, P<0.01 vs. control SAL in each case (2-way ANOVA). (C) Western blot analysis of C/EBPα protein levels in EWAT of control and mutant mice treated with salicylate (SAL) or vehicle (VEH). Representative blot show results of 3 animals per group. (D) Histogram shows quantification of C/EBPα levels normalized to TBP (TATA binding protein) and expressed as mean ± SEM. N=6 mice per group. *, P<0.05; **, P<0.01 (between genotypes) 2-way ANOVA). (E, F) Assessment of lipolysis as plasma glycerol (E) or free-fatty acids (FFAs) (F) concentration (μM) normalized to fat mass in control and mutant mice treated with salicylate (SAL) or vehicle (VEH) under basal conditions. Shown are mean values ± SEM. N=6 mice per group. *, P<0.05 vs. control SAL (2-way ANOVA). (G-L) Q-PCR determination of *IL-6* (G), *Tnfa* (H), *Mcp1* (I), *F4/80* (J), *Mrc1* (K) and *Clec10a* (L) mRNA expression in EWAT and SWAT of control and mutant mice treated with salicylate (SAL) or vehicle (VEH). Values were normalized to mRNA levels in VEH control mice, and are presented as mean ± SEM. N=6 mice per group. *, P<0.05; **, P<0.01 vs. control (within same treatment), or between treatments, as indicated (2-way ANOVA).

## Discussion

Our previous studies on ALK7 showed that deletion or inhibition of this receptor in otherwise healthy adipose tissue of lean mice prevents diet-induced catecholamine resistance, thereby enhancing energy expenditure and reducing weight gain when animals are switched to a high fat diet (Andersson et al., 2008; Guo et al., 2014). In the present study, we asked whether deletion of ALK7 in adipose tissue of mice that were already obese could similarly counteract further weight gain under a continued high caloric intake. We found that the effects were rather modest, paralleled by moderate changes in *Adrb* expression, beta-adrenergic signaling and lipolysis in the mutants. The contrast with our previous studies in lean mice suggested that, in addition to ALK7 activity, other factors contribute to suppress catecholamine signaling in adipose tissue in the obese state. Indeed, we observed that ALK7 deletion was much more effective when obese mice were switched to a low-caloric standard diet, reflected by a robust increase in *Adrb* expression, beta-adrenergic signaling, lipolysis, and energy expenditure in the mutants. Interestingly, diet switch led to a significant reduction in the expression of a series of inflammation markers in both control and mutant mice. This suggested that the overt inflammation state that characterizes adipose tissue in obesity may make adipocytes refractory to the effects of ALK7 deletion on adrenergic signaling and energy expenditure. Direct support for this notion was obtained from studies using Na-salicylate. As expected, salicylate treatment reduced the expression of inflammation markers in adipose tissue of obese mice, although it did not have any effect on adrenergic signaling or lipolysis. On the other hand, it synergized very strongly with ALK7 deletion, resulting in enhanced *Adrb* expression, increased lipolysis and reduced adipose tissue mass and body weight gain, even under sustained high caloric intake. We conclude from these observations that ALK7 signaling and inflammation work together to suppress β-adrenergic signaling in adipose tissue of obese mice, preserving fat mass. Both these pathways need to be inhibited to relieve this suppression, and thereby enhance lipolysis and energy expenditure.

In rodents as well as humans, obesity is associated with catecholamine resistance in adipose tissue, manifested as blunted β-adrenergic-mediated lipolysis and lipid oxidation (Arner, 1999; Jocken et al., 2008). High fat diet and obesity increase adipose tissue expression of activin B and GDF-3, two of the main ligands of ALK7 (Andersson et al., 2008; Sjöholm et al., 2006; Witthuhn and Bernlohr, 2001; Zaragosi et al., 2010), indicating enhanced ALK7 signaling in obesity. Lean mutant mice lacking ALK7 from birth, either globally or only in adipose tissue, gain less weight than wild type controls during diet-induced obesity, paralleled by increased *Adrb3* gene expression, adrenergic signaling, lipolysis and energy expenditure (Guo et al., 2014). One mechanism by which ALK7 suppresses catecholamine sensitivity upon nutrient overload is by limiting the expression of C/EBPα (this study and Lee et al., in preparation), a key determinant for transcription of the *Adrb3* gene (Dixon et al., 2001), which encodes the main β-adrenergic receptor responsible for catecholamine-induced lipolysis in adipose tissue of rodents. It has previously been shown that Smad3, a key intracellular mediator of signaling by TGF-β family receptors, including ALK7, interacts directly with C/EBPβ and C/EBPδ, thereby blocking their ability to induce *Cebpa* gene transcription (Choy and Derynck, 2003). By inhibiting C/EBPα expression, TGF-β has been shown to block differentiation of pre-adipocytes during the early stages of adipogenesis *in vitro* (Choy et al., 2000; Choy and Derynck, 2003). ALK7 does not affect adipocyte differentiation as it is only expressed in fully differentiated adipocytes (Guo et al., 2014; Kogame et al., 2006; Yogosawa et al., 2013), but can use a similar mechanism to suppress C/EBPα expression, thereby limiting β-adrenergic receptor levels and catecholamine sensitivity in mature adipocytes. The question then becomes why adipose tissue of obese mice is refractory to these effects and how this pathway intersects with inflammation signaling.

In obesity, constraints on adipocyte expansion and adipose tissue growth create stress on adipocytes, which respond by initiation of an inflammation program. Potential triggers of obesity-associated adipose tissue inflammation include adipocyte death, hypoxia and mechanical stress arising from interactions between expanding adipocytes and the extracellular matrix (Reilly and Saltiel, 2017). Although the effects of inflammation on insulin resistance in adipocytes and other tissues have been thoroughly investigated (reviewed in (Chawla et al., 2011; Hotamisligil, 2006; Reilly and Saltiel, 2017)), whether and how inflammation pathways affect catecholamine resistance and lipolysis in the obese state is less well understood. A study by Mowers et al. has pointed to two non-canonical members of the IKK kinase family, namely IKK∊ and TBK1, as potential links between inflammation signaling and catecholamine resistance in obesity, by their ability to desensitize lipolytic signaling (Mowers et al., 2013). Specifically, this study found that NF-κB signaling in response to inflammatory cytokines, such as TNF-α, increased expression of both IKK∊ and TBK1 kinases. These in turn phosphorylated and activated the major adipocyte phosphodiesterase PDE3B, thereby antagonizing cAMP signaling induced by β-adrenergic receptor activity (Mowers et al., 2013). In our experiments, we found that salicylate treatment reduced the expression of inflammatory cytokine genes *IL-6* and *Tnfa* but did not significantly affect adrenergic signaling on its own, as assessed by C/EBPα levels or lipolysis. We hypothesize that, in the presence of ALK7, reduction of phosphodiesterase activity may have limited effects on catecholamine signaling due to low levels of β-adrenergic receptors. Likewise, enhanced *Adrb3* gene expression following ALK7 deletion may not lead to increased lipolytic signaling in the obese state due to high phosphodiesterase activity induced by inflammatory cytokines. Our results suggest that only when both inflammation and ALK7 signaling are suppressed can catecholamine sensitivity be restored in adipose tissue of mice with pre-existing obesity.

Finally, our study has implications for the development of therapeutic strategies in human obesity and warrant efforts to identify safe and specific ALK7 inhibitors. ALK7 is not essential for life (Jörnvall et al., 2004), making it an attractive drug target. Life-style interventions may have a longer lasting impact if combined with suppression of ALK7 signaling. Likewise, safe, over-the-counter anti-inflammatory treatments, such as salicylate, may realize their full potential as anti-obesity therapeutics when administered along with an ALK7 inhibitor. The kinase domain of ALK7 is nearly identical to that present in related receptors for TGF-βs and activins, such as ALK5 and ALK4, respectively, and all current inhibitors targeting these kinases have broad effects on all three receptors (Byfield et al., 2004; Huynh et al., 2019; Inman et al., 2002). As ALK4 and ALK5 are more broadly expressed than ALK7 and involved in many different developmental and physiological processes (Gordon and Blobe, 2008; Morikawa et al., 2016), it will be necessary to target other domains in the ALK7 molecule to identify specific inhibitors with low toxicity. In this regard, decoy receptors and blocking antibodies may also offer useful avenues for development of ALK7 inhibitors. Regardless of these limitations, blockade of ALK7, if achieved with low toxicity and minimal side-effects, represents an important avenue to develop new strategies to combat human obesity and metabolic syndrome.

## Methods

### Animals

*Alk7*^flox/flox^ mice (Guo et al., 2014) were bred to *AdipoQ*^CreERT2^ mice (Sassmann et al., 2010) to generate *Alk7*^flox/flox^::*AdipoQ*^CreERT2^ mice, referred to as mutant mice throughout out the study. Mice were housed at 22±1°C and 45% humidity with ad libitum access to either standard Chow or high-fat diet (SD or HFD, respectively) as indicated, and drinking water. The HFD used contains 60% of calories from fat (Research Diet Inc, USA; Cat no. D12492). Daily food intake was assessed three times per week, starting two weeks after tamoxifen injection and for a period of four weeks. For the groups undergoing double diet switch, food intake was assessed during four weeks before the second switch, then during four weeks after the second switch. Cre recombinase-mediated *Alk7* deletion was induced by three daily consecutive injections of 25 mg/kg tamoxifen (Sigma, USA cat no. T5648) dissolved in saline solution containing 5% DMSO and 5% Tween-80. Body weight was recorded 3 or 4 times a week, as indicated. Na-salicylate (Sigma-Aldrich, Cat no. S3007) was administered in drinking water at 2g/L; control group received normal drinking water (vehicle). The mice were sacrificed after 9 weeks of this treatment. All experiments utilized male mice and were performed according to protocols approved by the IACUC of the National University of Singapore.

### Indirect calorimetry

Indirect calorimetry, food intake, and locomotor activity were assessed using a comprehensive lab animal monitoring system (Columbus Instruments) as previously described (Guo et al., 2014). Mice subjected to diet-switch were housed in metabolic cages individually with ad libitum access to standard diet or high fat diet, accordingly, and water. Mice were acclimatized to the cages for 24hs prior to a 48hs period of automated recordings. Oxygen consumption (VO_2_) and CO2 production (VCO_2_) were assessed through an open-circuit Oxymax. Ambulatory locomotor activity (XAMB) was measured by consecutive beam breaks in adjacent beams under 48hs period and presented as counts/min. Energy expenditure (EE) was calculated according to the formula EE = 3.815 × VO_2_ + 1.232 × VCO_2_ as previously described (Lusk, 1924).

### Measurements of plasma leptin and insulin, glucose and insulin tolerance tests

Mice were fasted for 6hs prior to collection of blood from the submandibular vein. Plasma was separated and kept frozen at −20°C until used. Plasma leptin and insulin levels were measured by ELISA using commercially available kits from ALPCO, USA (Leptin: Cat no. 22-LEPMS-E01; Insulin: Cat no. 80-INSMS-E01). For glucose tolerance test (GTT), mice were fasted for 6 hs, injected intraperitoneally with glucose (2g/kg, Sigma-Aldrich G8270), and blood glucose measured at 0, 15, 30, 60, 120 min post-injection from tail vein blood using a glucometer (FreeStyle, Abott). For insulin tolerance test (ITT), mice were fasted for 4 hs, injected intraperitoneally with insulin (0.75 U/Kg; Humulin R, Lilly), and glucose measured at 0, 15, 30, 60, 120 min post-injection from tail vein blood by ELISA using a commercial kit as described above.

### Basal and CL316243-induced lipolysis

Lipolysis was assessed as the levels of free glycerol and fatty acids in plasma normalized to fat mass (calculated as the combined weights of epididymal, retroperitoneal, perirenal and inguinal adipose tissue depots). For analysis of catecholamine-induced lipolysis, the β3-adrenergic receptor agonist CL316243 (Sigma-Aldrich, Cat no. C5976) was acutely administered by intraperitoneal injection at 1mg/kg body weight. Blood was collected 30 min after the injection and plasma was separated for assessment of glycerol and free fatty acids. Glycerol and free-fatty acids were measured using commercial kits Free Glycerol Reagent (Sigma, USA; Cat. No: F6428) and Half-micro test (Roche, Germany; Cat. No. 11 383 175 001), respectively, according to manufacturers’ instructions.

### Histology and TUNEL

Epididymal and inguinal white adipose tissues were collected and fixed in 4% paraformaldehyde for 48hs at 4°C. The tissues were then dehydrated, cleared in xylene, embedded in paraffin wax and sectioned at 5 μm. Hematoxylin and eosin staining was performed as per standard protocols. The stained slides were imaged by wide field microscopy (Leica Microsystems, Germany), and images were analyzed to quantify adipocyte size by Image J software (NIH, USA). At least four fields were randomly selected for analysis from each tissue section per mouse. TUNEL staining was performed using a protocol modified from the ApopTag peroxidase in situ apoptosis detection kit (Merck-Millipore, Cat no. S7100). Paraffin sections of adipose tissues were prepared as above, then de-paraffinized and treated with 200 μg/ml Proteinase K for 15 mins. Digoxigenin-labelled nucleotides were enzymatically added to free 3’-OH DNA termini by terminal deoxynucleotidyl transferase as per manufacturer’s instructions. The incorporated digoxigenin-labelled nucleotides were detected by anti-digoxigenin-POD conjugated antibodies (Dilution 1.5 IU/ml; Roche, Cat no. 11633716001). Fluorescent signals were obtained by peroxidase-mediated Alexa Fluor 488 tyramide labelling (Alexa fluor 488 Tyramide SuperBoost kit, Thermofisher Scientific, Cat no. B40922). Slides were counterstained with DAPI and mounted in ProLong Diamond antifade mounting medium (Thermofisher Scientific, Cat no. P36970). Images were captured in a confocal microscope (Leica Microsystems, Germany) and analyzed by Image J software (NIH, USA).

### Real-time PCR

Adipose tissues were homogenized in TRIzol (Invitrogen, Cat. No. 15596026), centrifuged at 15,000 rpm to remove debris and the supernatant was collected. 200 μL chloroform was added per milliliter of TRIzol, mixed by vortexing and centrifuged at 15,000 rpm for 15 min. The aqueous phase was collected, an equal amount of ethanol was added and loaded on to the kit column and RNA further purified and cleaned using RNeasy kit (QIAGEN, USA) following manufacturer’s instructions. The organic phase was saved for subsequent immunoblotting analysis of proteins (see below). 500ng RNA was used to synthesize cDNA using High capacity cDNA reverse transcription kit (Applied Biosystems, Cat no. 4390778) as per instructions provided in the kit. Quantitative, real-time PCR was run in the reaction plate containing diluted cDNA, SYBR select master mix (Thermo Fisher Scientific, cat no. 4472919), and forward and reverse primers (Supplementary File 1). To quantify gene expression, the CT value for the gene of interest was normalized to that of mRNA encoding TATA-binding protein (TBP).

### Immunoblotting

For immunoblotting analysis, proteins were precipitated from the organic phase obtained with the TRIzol reagent (see above) as previously described (Srivastava et al., 2020). Protein pellets were dissolved in sample buffer, supplemented with β-mercaptoethanol and heated at 95°C for 10 min. The samples were centrifuged at 10,000 rpm for 10 min to remove particulates, and protein content was estimated using Pierce 660nm protein assay reagent (Thermo Fisher Scientific, Cat no. 22660). Equal amount of protein was loaded onto SDS-PAGE gels for separation. The separated proteins were electro-transferred to PVDF membranes (GE healthcare, Cat no. 10600021). These were subsequently blocked with 5% milk in Tris-buffered saline with 0.1% Tween 20 (TBST) and incubated overnight with primary antibodies (Supplementary File 2). The blots were washed and immunoreactive bands were detected by incubation with anti-rabbit HRP conjugated antibody (dilution 1:2000 in blocking solution; Cell signaling, Cat no. 7074S) for 1h at room temperature. The blots were then washed three times with TBST, and developed with SuperSignal West Femto maximum sensitivity substrate (Thermo Fisher Scientific, Cat no. 34095). Quantification was done with Image J software (NIH, USA) after exposing to X-ray films. GAPDH or TBP were used as reference proteins for normalization.

### Statistical analysis

Statistical analyses were performed by using Prism 5 software (Graph Pad, SPSS IBM Corporation). The results are presented as mean ± Standard error of the mean (SEM). One-way ANOVA or Two-way AVOVA were used to test statistical significance as per the requirement of the experiments. The level of statistical significance was set at P<0.05 for all the analyses (*). All P values are reported in the figure legends.

## Acknowledgements

We thank Goh Ket Yin for technical assistance and Meng Xie for comments on the manuscript. This research was funded by grants NMRC/CBRG/0107/2016 from the Singapore National Medical Research Council, Aspiration Fund Partner from the National University of Singapore, and 2016-01538 from the Swedish Research Council (Vetenskapsrådet).

---

**Supplementary Figure 1.**
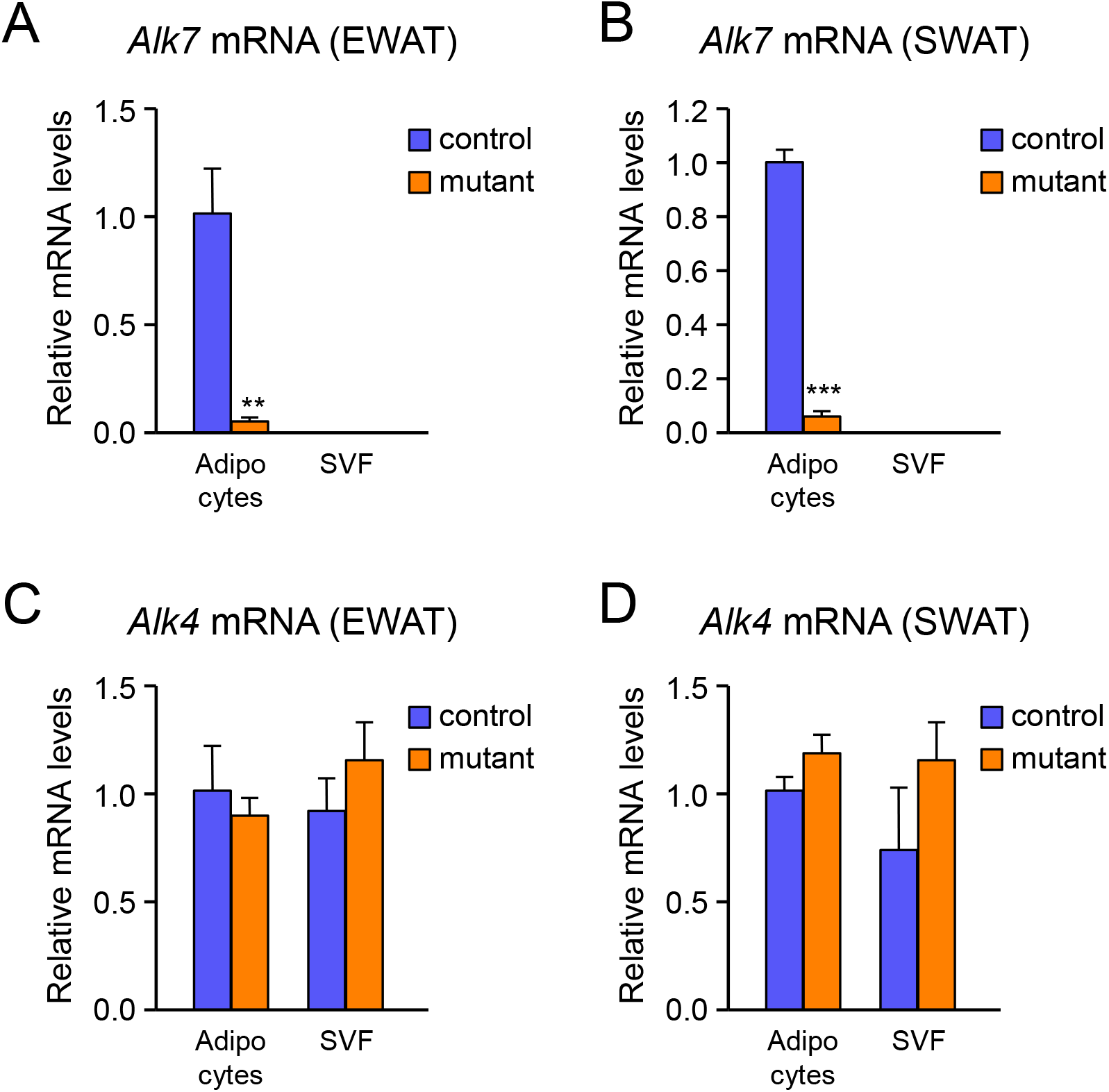
Conditional ablation of ALK7 expression in adipose tissue of obese mice. Relative expression of *Alk7* (A, B) and *Alk4* (C, D) mRNAs in adipocytes and stromal vascular fraction (SVF) of EWAT (A, C) and SWAT (B, D) two weeks after tamoxifen-induced deletion in *Alk7*^flox/flox^ (control) and *Alk7*^flox/flox^::*AdipoQ*^CreERT2^ (mutant) mice kept for 12 weeks on high-fat diet. *Alk7* mRNA was undetectable in SVF samples. ***, P<0.001 vs. control (2-way ANOVA).

**Supplementary Figure 2.**
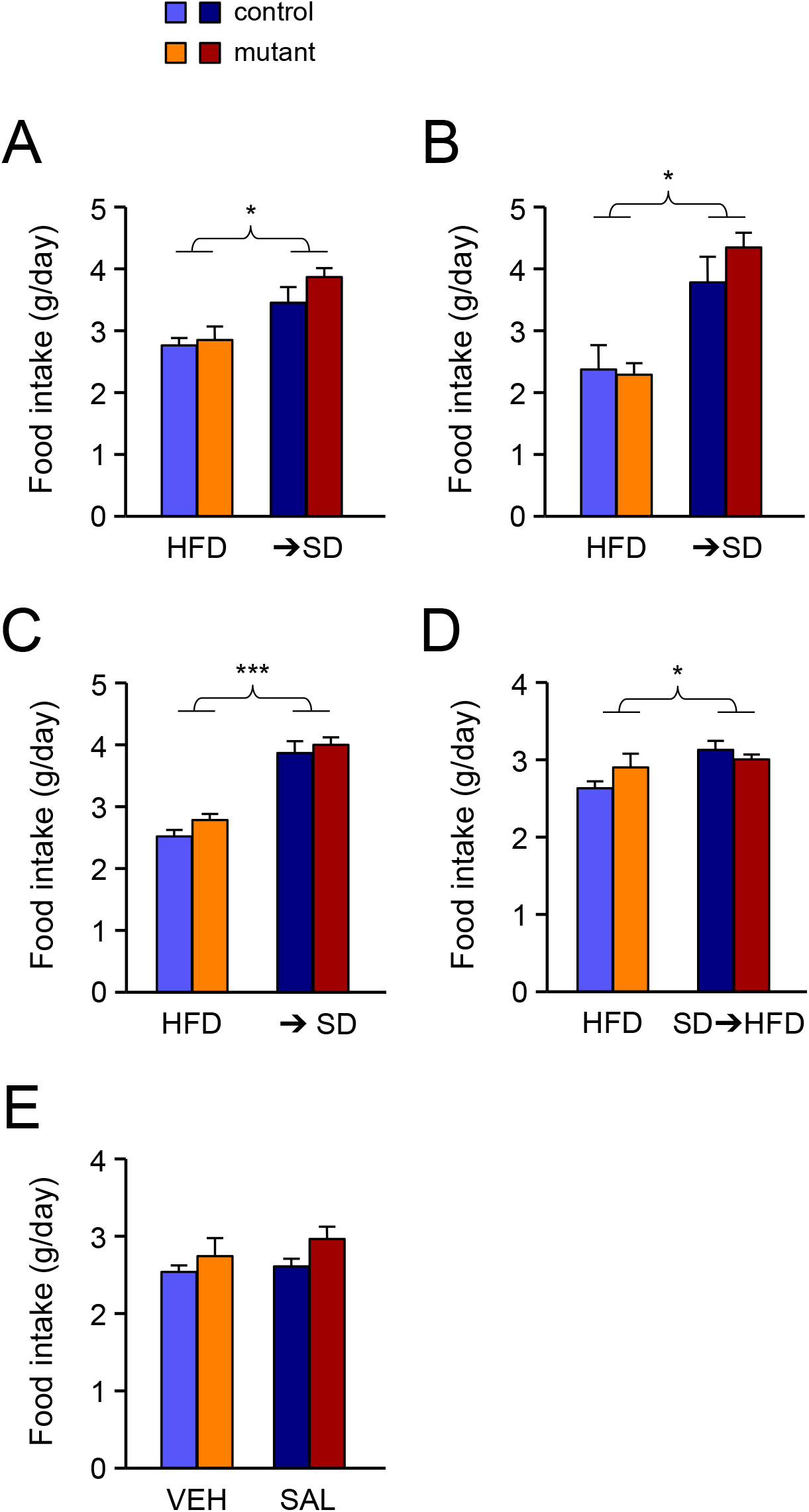
Food intake in control and mutant mice subjected to diet switch or salicylate treatment. Average (±SEM) daily food intake expressed in grams. (A) Single diet switch (➔SD) and constant HFD cohorts in normal cages. (B) Single diet switch (➔SD) and constant HFD cohorts in CLAMS metabolic cages. (C) Double diet switch (➔SD) and constant HFD cohorts prior to second switch. (D) Double diet switch (SD➔HFD) and constant HFD cohorts after second switch. (E) Salicylate and vehicle cohorts. *, P<0.05; ***, P<0.001 between diets or treatments, as indicated. N=6 mice per group.

**Supplementary Figure 3.**
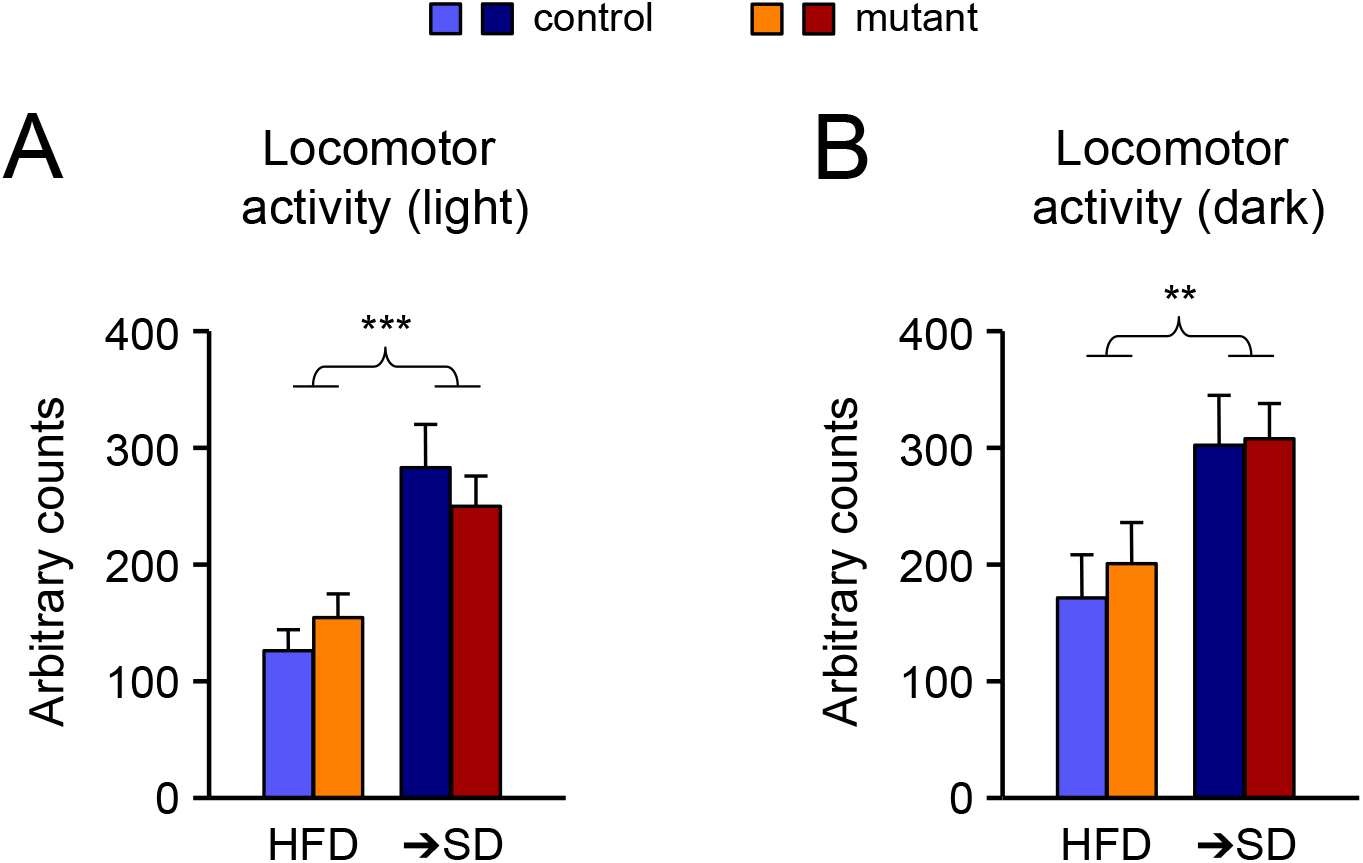
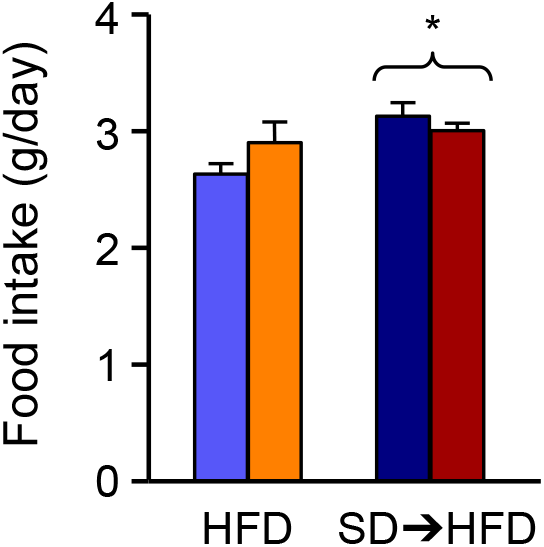
Locomotor activity in control and mutant mice subjected to diet switch. Average (±SEM) locomotor activity expressed in arbitrary counts during light (A) or dark (B) periods. **, P<0.01; ***, P<0.001 between diets, as indicated. There was no significant difference between genotypes. N=6 mice per group. (2-way ANOVA).

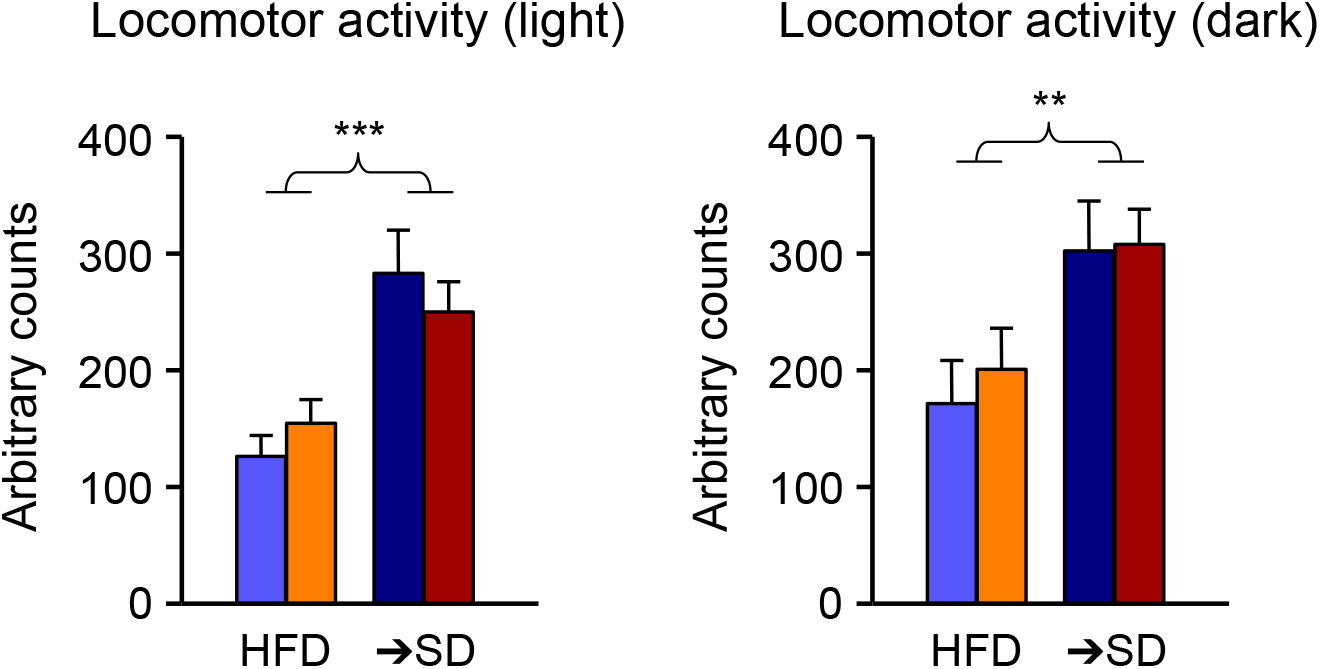

**Supplementary File 1:**
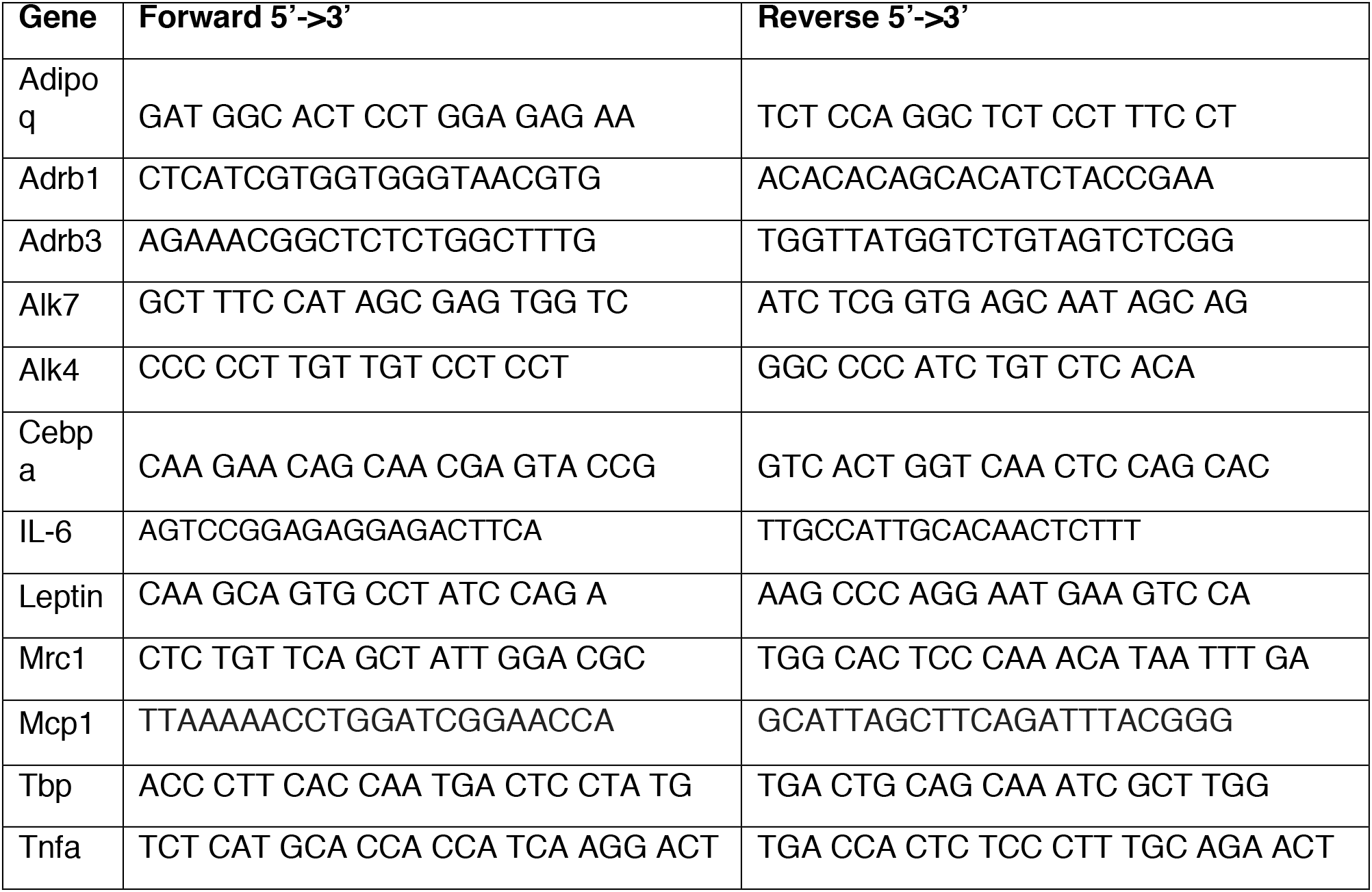
PCR primers.

**Supplementary File 2:**
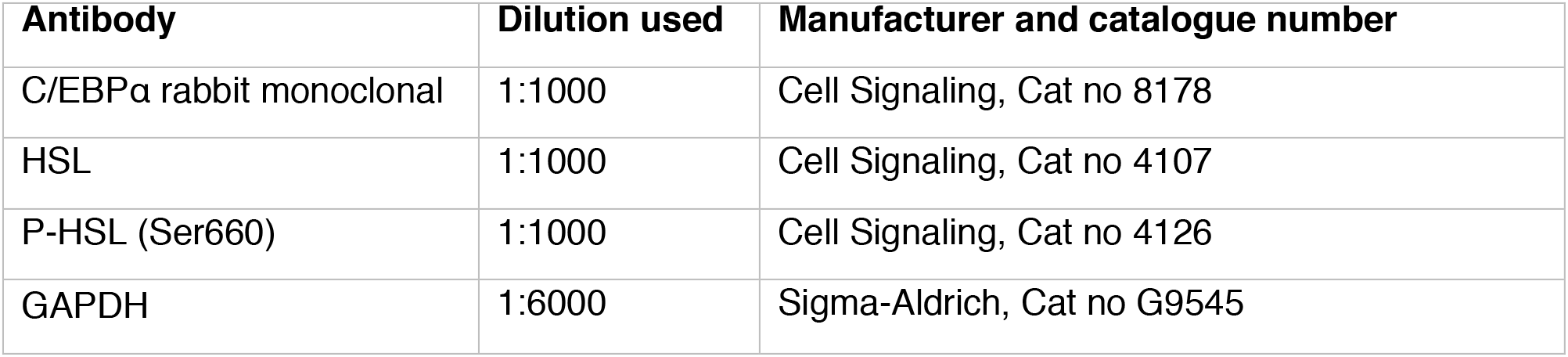
Primary antibodies.

| Antibody | Dilution used | Manufacturer and catalogue number |
| --- | --- | --- |
| C/EBPα rabbit monoclonal | 1:1000 | Cell Signaling, Cat no 8178 |
| HSL | 1:1000 | Cell Signaling, Cat no 4107 |
| P-HSL (Ser660) | 1:1000 | Cell Signaling, Cat no 4126 |
| GAPDH | 1:6000 | Sigma-Aldrich, Cat no G9545 |

